# A critical re-evaluation of fMRI signatures of motor sequence learning

**DOI:** 10.1101/2020.01.08.899229

**Authors:** Eva Berlot, Nicola J. Popp, Jörn Diedrichsen

## Abstract

Despite numerous studies, there is little agreement about what brain changes accompany motor sequence learning, partly because of a general publication bias that favors novel results. We therefore decided to systematically reinvestigate proposed functional magnetic resonance imaging correlates of motor learning in a preregistered longitudinal study with four scanning sessions over 5 weeks of training. Activation decreased more for trained than untrained sequences in premotor and parietal areas, without any evidence of learning-related activation increases. Premotor and parietal regions also exhibited changes in the fine-grained, sequence-specific activation patterns early in learning, which stabilized later. No changes were observed in the primary motor cortex (M1). Overall, our study provides evidence that human motor sequence learning occurs outside of M1. Furthermore, it shows that we cannot expect to find activity increases as an indicator for learning, making subtle changes in activity patterns across weeks the most promising fMRI correlate of training-induced plasticity.

## Introduction

Humans have the remarkable ability to learn complex sequences of movements. While behavioural improvements in sequence learning tasks are easily observable, the underlying neural processes remain elusive. Understanding the neural underpinnings of motor sequence learning could provide clues about more general mechanisms of plasticity in the brain. This motivation has led numerous functional magnetic resonance imaging (fMRI) studies to investigate the brain changes related to motor sequence learning. However, there is little agreement about how and where in the brain learning-related changes are observable. Previous studies include reports of signal increases across various brain regions (Floyer-Lea & Matthews, 2005; Grafton, Hazeltine, & Ivry, 1995; Hazeltine, Grafton, & Ivry, 1997; Karni et al., 1995; Lehéricy et al., 2005; Penhune & Doyon, 2002), as well as signal decreases (Jenkins, Brooks, Nixon, Frackowiak, & Passingham, 1994; Peters, Lee, Hedrick, Neil, & Komiyama, 2017; Toni, Krams, Turner, & Passingham, 1998; Ungerleider, Doyon, & Karni, 2002; Wiestler & Diedrichsen, 2013), nonlinear changes in activation (Ma et al., 2010; Xiong et al., 2009), spatial shifts in activity (Lehéricy et al., 2006; Steele & Penhune, 2010), changes in multivariate patterns (Wiestler & Diedrichsen, 2013; Wymbs & Grafton, 2015), and changes in inter-regional functional connectivity (Bassett, Yang, Wymbs, & Grafton, 2015; Bassett et al., 2010; Doyon et al., 2002; Mattar et al., 2016). Additionally, some experiments have matched the speed of performance (Karni et al., 1995; Penhune & Doyon, 2002; Steele & Penhune, 2010; Lehéricy et al., 2005; Seidler et al., 2002, 2005), while others have not (Bassett et al., 2015; Lutz, Koeneke, Wüstenberg, & Jäncke, 2004; Wiestler & Diedrichsen, 2013; Wymbs & Grafton, 2015). Given that fMRI analysis has many degrees of freedom, these inconsistencies may not be too surprising. However, the implicit pressure in the publication system to report findings may also have contributed to a lack of coherency. To address this issue, we designed a comprehensive longitudinal study of motor sequence learning that allowed us to systematically reinvestigate previous findings. In order to increase transparency, we pre-registered the design, as well as all tested hypotheses on the Open Science Framework (Berlot, Popp, & Diedrichsen, 2017; https://osf.io/etnqc), and make the full dataset available to the research community.

The main aim of our study was to systematically evaluate different ideas of how learning-related changes are reflected in the fMRI signal. In the context of motor sequence learning, the most commonly examined brain region is the primary motor cortex (M1). Previous reports of increased M1 activation after long-term learning have been interpreted as additional recruitment of neuronal resources for trained behavior, taken to suggest the skill is represented in M1 (Floyer-Lea & Matthews, 2005; Karni et al., 1995, 1998; Lehéricy et al., 2005; Penhune & Doyon, 2002; for a review see Dayan & Cohen, 2011; Fig. 1a). Since then, several pieces of evidence have suggested that sequence-specific memory may not reside in M1 (Beukema, Diedrichsen, & Verstynen, 2019; Wiestler & Diedrichsen, 2013; Yokoi & Diedrichsen, 2019). However, some of these reports studied skill acquisition over a course of a few days, while human skill typically evolves over weeks (and months) of practice. Therefore, including several weeks of practice, might be more suitable to test whether, and at what time point, M1 develops skill-specific representations.

**Figure 1.**
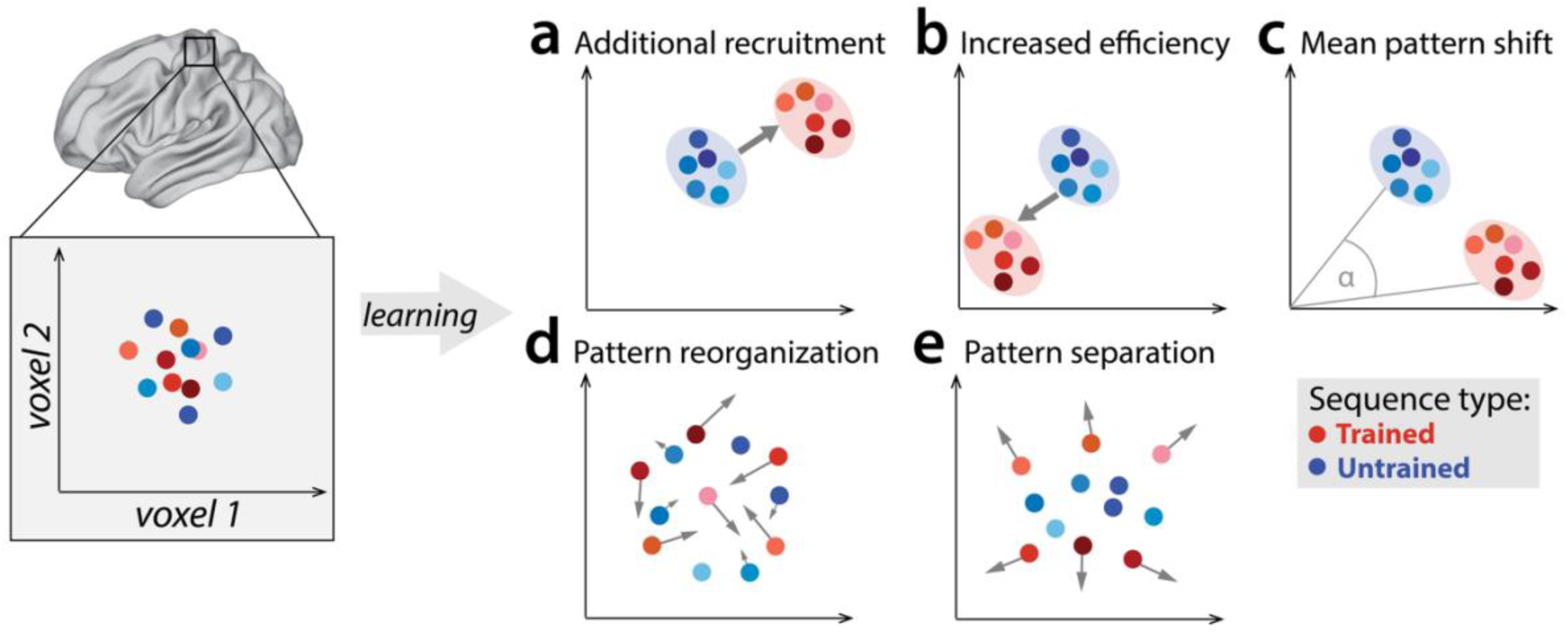
Potential fMRI signatures of learning in a specific brain area. Each panel shows hypothetical activation for the six trained sequences (red) and the six untrained sequences (blue) in the space of two hypothetical voxels. **a)** Activation could increase during learning across voxels, indicating additional recruitment of resources involved in skilled behavior. **b)** Activation could decrease across voxels, implying that the region performs its function more efficiently. **c)** Some voxels (x-axis) could increase activation with training, while others (y-axis) could decrease. This would lead to a shift of the overall activity pattern in the region without an overall net change in activation. **d)** Activation patterns specific to each trained sequence could undergo more change than untrained sequences, reflective of plastic reorganization of the sequence representation. Arrow length in the figure indicates the amount of reorganization. **e)** One specific form of such reorganization would be increasing dissimilarities (pattern separation) between activity patterns for individual trained sequences.

Outside of M1, learning-related activation changes have been reported in premotor and parietal areas (Grafton, Hazeltine, & Ivry, 2002; Hardwick, Rottschy, Miall, & Eickhoff, 2013; Honda et al., 1998; Penhune & Doyon, 2002; Tamás Kincses et al., 2008; Vahdat et al., 2015), with activation increases commonly interpreted as increased involvement of these areas in the skilled behavior. Yet, recent studies have mostly found that, as the motor skill develops, activation in these areas predominantly decreases (Penhune & Steele, 2012; Wiestler & Diedrichsen, 2013; Wu et al., 2004). Such reductions are harder to interpret as they could reflect a reduced areal involvement in skilled performance or, alternatively, more energy efficient implementation of the same function (Fig. 1b) (Picard, Matsuzaka, & Strick, 2013; Poldrack et al., 2005). To complicate things further, regional activity increases and decreases could occur simultaneously in the same area (Fig. 1c; Steele & Penhune, 2010). In such a scenario, the net activation in the region would not change, yet, the trained sequences would engage slightly different subpopulations of the region than untrained sequences.

A variant of this idea is that each specific sequence becomes associated with dedicated neuronal subpopulation (and hence fMRI activity pattern). Such a representation would form the neural correlate of sequence-specific learning – the part of the skill that does not generalize to novel, untrained motor sequences (Karni et al., 1995). Sequence-specific activation patterns should change early in learning (Fig. 1d), when behavior improves most rapidly, and stabilize later, once the skill has consolidated and an optimal pattern is established (Peters et al., 2017). One possible way in which sequence-specific patterns could reorganize is by becoming more distinct from one another (Fig. 1e; Wiestler & Diedrichsen, 2013). Having a distinctive code for each sequence might be of particular importance to the system in a trained state, allowing it to produce different dynamical sequences, while avoiding confusion or “tangling” of the different neural trajectories (Russo et al., 2018).

To systematically examine the cortical changes associated with motor sequence learning, we carried out a longitudinal study over 5 weeks of training with 4 sessions of high-field (7 Tesla) fMRI scans. Behavioural performance in the first three scanning sessions was imposed to the same speed of performance. This allowed us to inspect whether examined fMRI metrics reflect brain reorganization, independent of behavioral change. However, controlling for speed incurs the danger of not tapping into neural resources that are necessary for skilled performance (Orban et al., 2010; Poldrack, 2000). We therefore compared the fMRI session with paced performance at the end of behavioural training with one acquired with full speed performance (Fig. 2). This manipulation allowed us to systematically assess the role of speed on the fMRI metrics of learning, thereby addressing an important methodological problem faced by virtually every study on motor learning.

**Figure 2.**
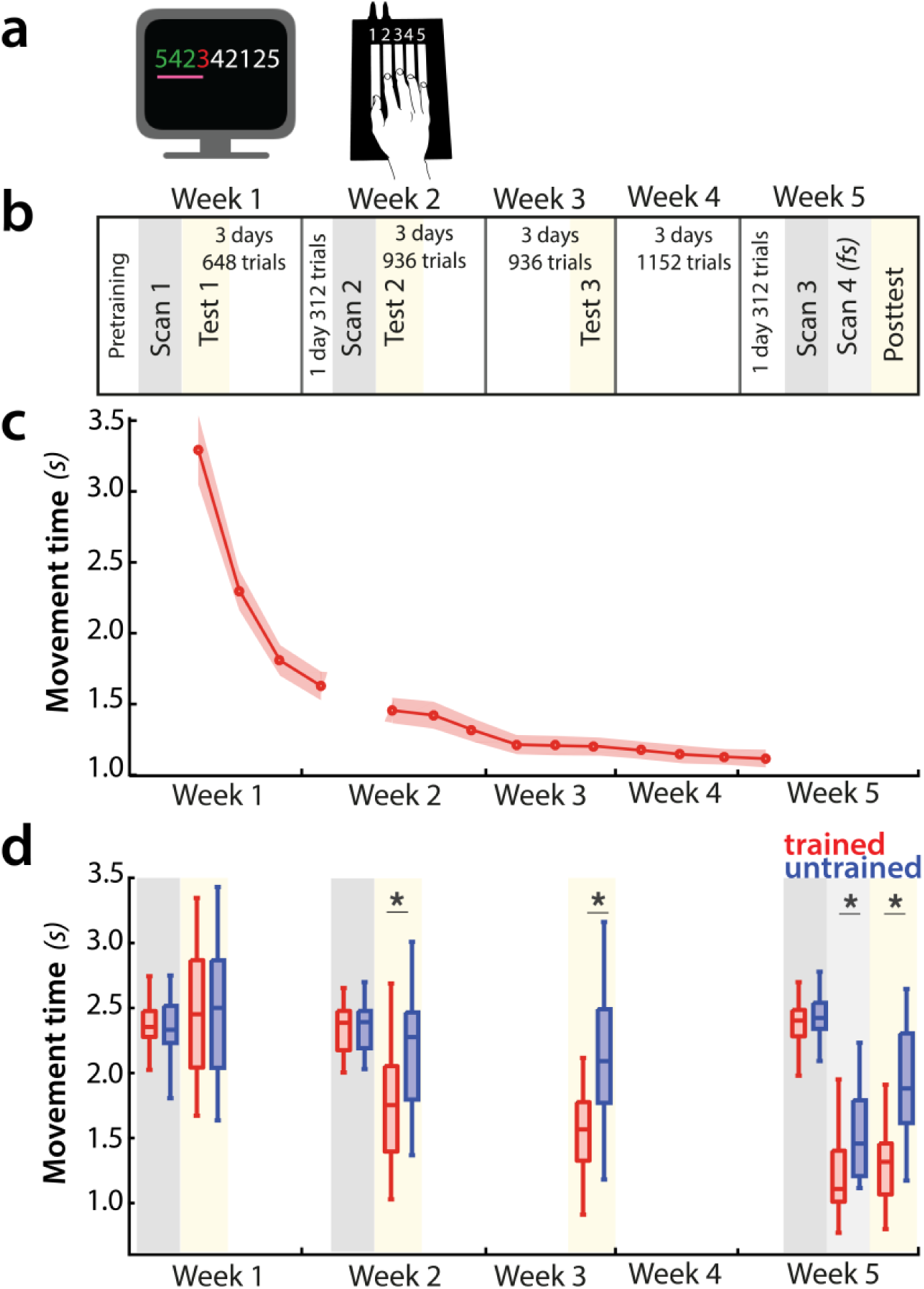
Experimental design and paradigm. **a)** Apparatus and task. Participants were trained to perform six 9-item sequences on a keyboard device. For each finger press, the corresponding digit on the screen turned green (correct) or red (incorrect). During fMRI scans 1-3, an expanding pink line under the numbers indicated the pace at which participants had to press the keys. See supplementary figure **S2** for trial structure during scanning sessions. **b)** Training protocol lasted for 5 weeks, and included four behavioral test sessions (yellow underlay) and four scans (grey underlay). Scans 1-3 were performed at a paced speed, while scan 4 performance was full speed *(fs)*. **c)** Average group performance executing trained sequences across the training sessions, measured in seconds. The average movement time (MT) decreased with learning. Shaded area denotes between-subject standard error. **d)** Performance during scanning sessions and behavioral tests, measured in seconds. Performance of trained sequences improved across all subsequent behavioral test sessions. Performance improved also for untrained sequences from week 2 onwards, suggesting some transfer in learning, but performance was still faster for trained sequences, indicating sequence-specific learning. Error bars indicate between-subject standard error. Stars denote significance levels lower than *p*<.001.

## Results

### Speed of sequence execution increases with learning

We trained 26 participants to perform six 9-digit sequences with their right hand on a keyboard device (Fig. 2a). During training, they received visual feedback (green for correct and red for incorrect presses) and were rewarded for both accuracy and speed (see Materials and Methods). Over the course of 5 weeks, participants practiced ~4000 trials (Fig. 2b). This led to substantial performance improvement, with the average movement time (MT) to complete a sequence decreasing from an initial 3.2 seconds to 1.2 seconds at the end of the training (Fig. 2c). The training regime was complemented with behavioral assessments on four occasions designed to specifically assess participants’ performance on trained sequences relative to untrained sequences (Fig. 2d, yellow underlay). Prior to training (test day 1), the speed of sequence execution did not differ between trained and untrained sequences. For all subsequent sessions, MTs were significantly faster for trained than untrained sequences (*p*<.001), implying sequence-specific learning. Additionally, performance of trained sequences improved between all subsequent sessions, even after week 3 (week 3-5: *t*_(25)_=5.49, *p*=1.1e-5). Thus, participants’ performance of trained sequences improved across the five weeks.

To assess fMRI changes with learning, participants underwent four fMRI scans (1st scan: before the main training; 2_nd_ scan: week 2; 3_rd_ & 4_th_ scan: week 5), performing both trained and untrained sequences (Fig. 2d – grey underlay). During the first three sessions, participants were paced with a metronome so that all sequences, trained and untrained, were performed at the same speed as in the first scan. Performance in the fourth session was at maximum speed, resulting in significantly lower MTs for trained compared to untrained sequences (Fig. 2d). To assess different neural signatures of observed behavioral learning, we first examined how the overall evoked activation changed over weeks of training for the same speed of movement.

### Overall activation does not change in M1

First, we re-investigated the classical finding that activity, measured as the percent BOLD signal change relative to rest, increased in M1 for matched performance after long-term training (Karni et al., 1995; Fig. 1a). Our task elicited activation in a range of cortical areas (Fig. 3a for session 1 – i.e., prior to learning). A region of interest (ROI) analysis of the hand area of M1, contralateral to the performing hand, however, showed no significant change across weeks (Fig. 3b, *F*_(2,50)_=1.82, *p*=.17). Neither did we find any difference between trained and untrained sequences (*F*_(1,25)_=0.19, *p*=.66), or a significant interaction between the two (*F*_(2,50)_=2.01, *p*=0.14).

**Figure 3.**
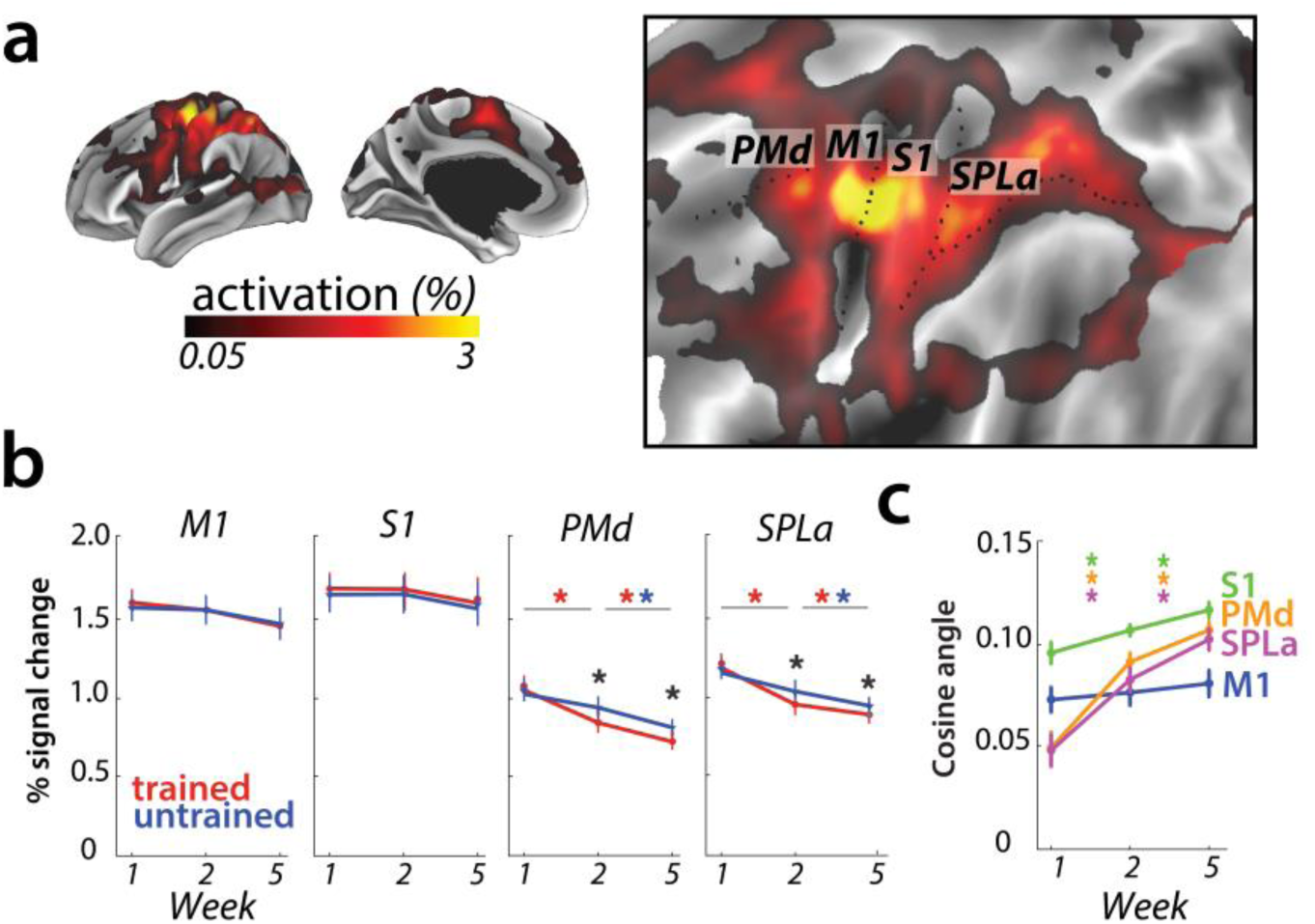
Overall activation and changes with learning in defined regions of interest. **a)** Average activation during production of any sequence in scanning session 1 (prior to learning) in the hemisphere contralateral to the performing hand. Activation was contrasted against resting baseline. On the right, activation map is presented on a flattened surface, corresponding to surface maps in other figures. **b)** Changes in activation across predefined areas – primary motor cortex (M1), primary somatosensory cortex (S1), premotor dorsal area (PMd) and superior parietal lobule – anterior (SPLa). No significant changes in activation were observed in M1 or S1 across weeks or between trained and untrained sequences (* indicates *p*<.01). Error bars indicate between-subject standard error. **c)** The cosine angle dissimilarity between average trained and untrained sequence across scanning weeks. The cosine angle increased significantly across weeks in PMd, SPLa and S1, but not M1 (* indicates *p*<.05). Error bars indicate between-subject standard error.

The absence of overall activity changes, however, should not be taken as evidence for an absence of plasticity in the region. It is possible that some subregions of M1 increased in activation for learned sequences, while other decreased, as suggested by Steele and Penhune (2010). Such mixed changes would result in a shift of the overall pattern, which would lead to an increase in the angle between the mean activity pattern for trained and untrained sequences (Fig. 1c). Because we calculated the angle between activity patterns for each participant separately, this criterion does not assume that the observed shift is spatially consistent across individuals – any idiosyncratic shift could be detected. Therefore it serves as a sensitive statistical criterion to detect shifts in spatial location of activation, which were previously reported only descriptively (Steele & Penhune, 2010).

However, in M1, the averaged cosine angle (Fig. 3c) remained unchanged across the weeks (*F*_(2,50)_=1.71, *p*=.19), indicating that the average activity pattern remained comparable across trained and untrained sequences. In sum, we found no evidence for activation increases (Karni et al., 1995), decreases, or relative shifts in activation patterns (Steele & Penhune, 2010) in M1.

### Learning-related activation changes in premotor and parietal areas

To investigate activation changes in areas outside of M1, we calculated changes in activity between the weeks in a map-wise approach (Fig. 4a). Over the three measurement time points, we found no reliable activation increases in any cortical area that was activated by the task in week 1. Instead, we observed widespread learning-related reductions in activity in premotor and parietal areas (Fig. 4a), in line with our pre-registered prediction. These activation reductions were observed across both subsequent sessions (i.e. weeks 1-2, weeks 2-5) for trained and untrained sequences, with bigger reductions for trained sequences. In weeks 2 and 5, trained sequences elicited overall lower activity than untrained sequences (Fig. 4b; see supplementary figure **S4** for statistical maps). These learning-related reductions in activity were also statistically significant in our predefined ROIs in premotor (dorsal premotor cortex – PMd) and parietal cortices (anterior superior parietal lobule – SPLa) (Fig. 3b): In a 3 (week) × 2 (sequence type) ANOVA on observed activation both main effects and interaction were highly significant in PMd (week: *F*_(2,50)_=17.47, *p*=1.77e-6; sequence type: *F*_(1,25)_=11.86, *p*=2.03e-3; interaction: *F*_(2,50)_=13.22, *p*=2.46e-5) as well as in SPLa (week: *F*_(2,50)_=19.14, *p*=6.73e-7; sequence type: *F*_(1,25)_=19.36, *p*=1.77e-4; interaction: *F*_(2,50)_=21.59, *p*=1.74e-7). In contrast, no main effect of week was observed in S1 (*F*_(2,50)_=0.44, *p*=.85). There was a significant main effect of sequence type (*F*_(1,25)_=6.32, *p*=.019), but none of the post-hoc t-tests revealed a significant difference. The week × sequence type interaction was not significant in S1 (*F*_(2,50)_=0.17, *p*=.84). Thus, we observed widespread activation decreases with learning across secondary and association cortical areas.

**Figure 4.**
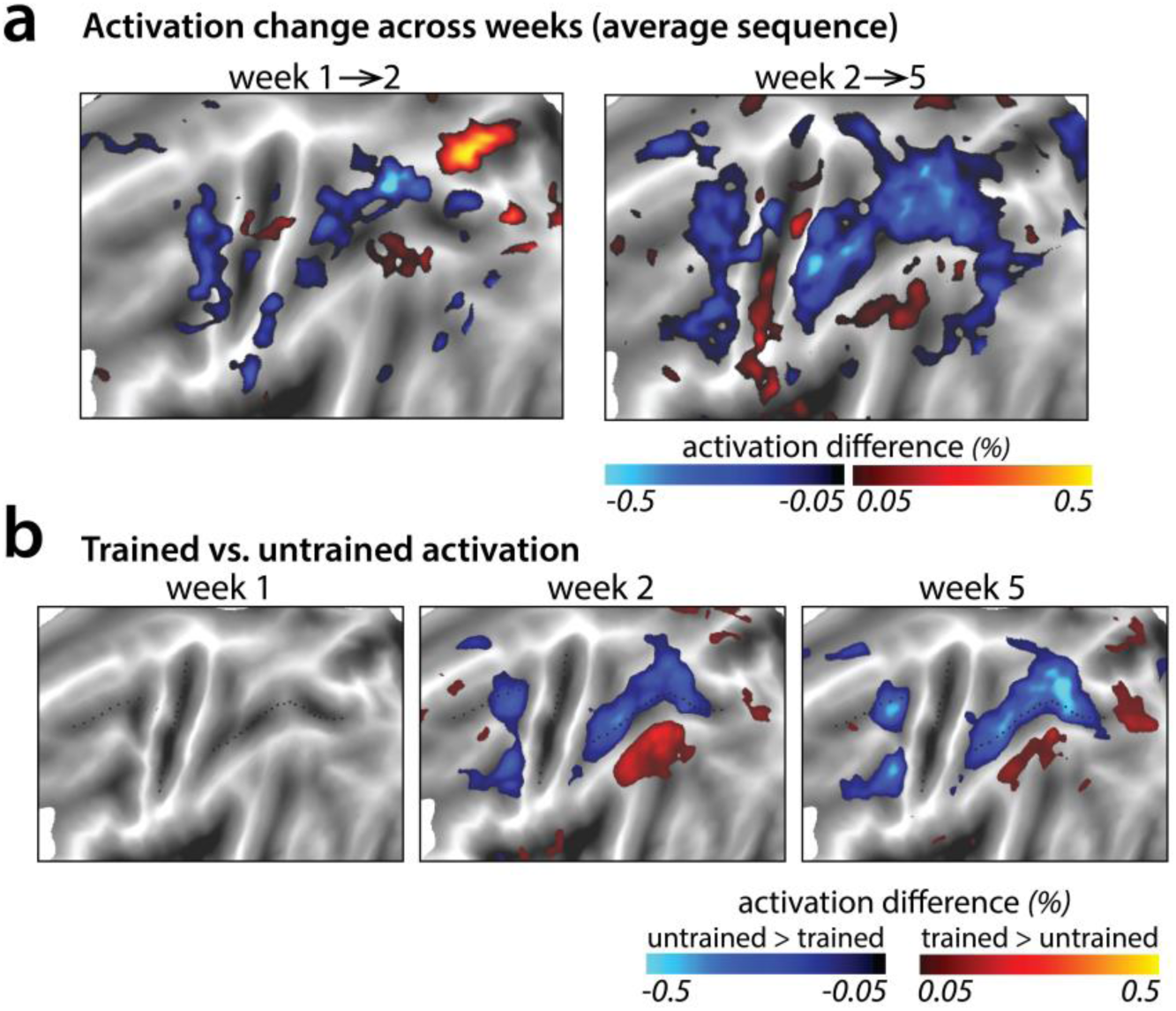
Changes in average activation across the cortical surface. **a)** Average change in activation across subsequent sessions. Activation was measured as difference in percent signal change relative to the resting baseline. Activation decreased (blue shades) in motor-related regions across sessions during sequence execution. **b)** Contrast of activation for trained vs. untrained sequences per scanning session. In weeks 2 and 5, trained sequences elicited lower activation in motor-related regions than untrained sequences (blue shades; see supplementary figure **S4** for *t*-maps and statistical quantification of activation clusters). Areas with observed increases in activation for trained sequences (red shades) lie in the default mode network that showed on average lower activity during task than rest.

In a few smaller areas, activation increased with learning (red patches in Fig. 4a-b). This was observed uniformly in areas with activity at or below baseline – thus these changes reflect decreased suppression of activity rather than increases. It is likely that these activity increases are not task relevant, but instead reflect the increasing automaticity and lower need for central attentional resources with learning (see Discussion).

We also examined whether there were, in addition to the overall activity decreases, shifts in the average activity patterns in the predefined regions of interest (Fig. 1c). As for M1, we calculated the cosine angle dissimilarity (see Materials and Methods) between the average activity patterns for trained and untrained sequences, separately for each scanning session. Figure **5a** shows cosine angle dissimilarities between trained and untrained sequences in PMd, displayed using multidimensional scaling (MDS). Patterns for trained sequences moved away from the starting point over weeks, and became more different from untrained patterns. Both in parietal and premotor areas there was clear evidence for a shift – cosine angular dissimilarity between the average trained and untrained sequence activation increased significantly across weeks (PMd: *F*_(2,50)_=23.63, *p*=5.98e-8; SPLa: *F*_(2,50)_=23.19, *p*=7.49e-8) (Fig. 3c). S1 also showed a significant increase in cosine dissimilarity between trained and untrained patterns with learning (*F*_(2,50)_=8.68, *p*=5.79e-4). These changes, however, were much less pronounced than those observed in premotor and parietal areas.

To investigate whether the observed changes in the overall activity patterns in premotor and parietal areas were spatially consistent across individuals, we normalized (z-scored) activation maps in each region and assessed the relative contribution of subregions to overall activation in weeks 1 and 5 (Fig. 5b). Comparing the pattern of activation revealed that before training (week 1, blue) sequences elicit relatively more activation in rostral parts of the premotor and supplementary motor areas, and that activity was more caudal after training (week 5, red; Fig. 5c displays the cross-section of relative activation changes). Some differences were also observed in the posterior parietal cortex, with activation shifting from more posterior to anterior subregions after learning (Fig. 5c). Altogether, these results show that with learning, the execution of sequences relies on slightly different subareas within premotor and parietal regions.

**Figure 5.**
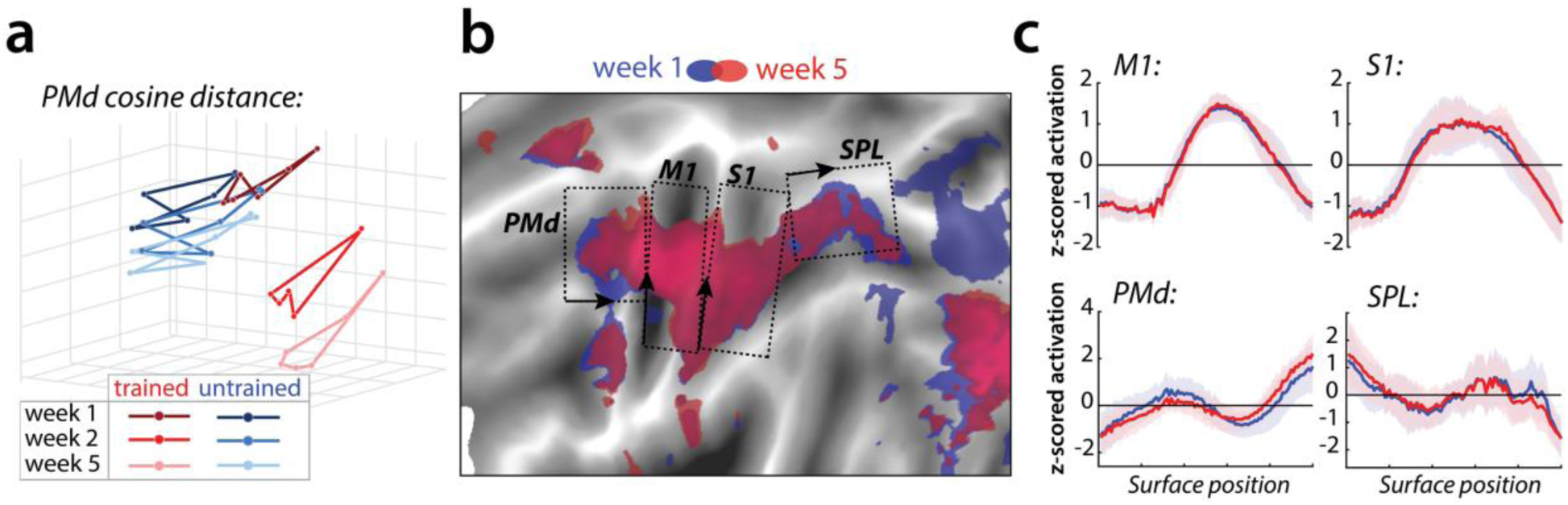
Relative change in evoked activation. **a)** Multidimensional scaling plot of cosine angle dissimilarities for trained and untrained sequences in premotor dorsal area (PMd) across weeks 1-5. Each dot represents a single sequence, and dots are connected for each session and sequence type separately. Trained sequences on average become more distant from untrained sequences with learning. Untrained sequences on average also progress across weeks, but less than trained sequences. **b)** Normalized activation plots for trained sequences in week 1 (blue) and 5 (red). The arrows and brackets indicate the direction and range of activation cross-sections presented in c). Areas: dorsal premotor cortex (PMd), primary motor cortex (M1), primary somatosensory cortex (S1), superior parietal lobule (SPL). **c)** Cross-section of elicited activation for trained sequences in defined areas, in weeks 1 (blue) and 5 (red).

### Sequence-specific activity patterns reorganize early in learning

Our analyses so far have been concerned with changes in the overall pattern of trained vs. untrained sequences, and showed widespread reductions in activation and some more subtle changes in relative location. The sequence-specific performance advantage, however, indicates that the brain must represent specific sequences – i.e. there should be activity patterns that are unique to each individual sequence. Sequence-specific learning should then be reflected in changes of these sequence-specific activity patterns with learning (Fig. 1d). Consistent with previous results (Wiestler & Diedrichsen, 2013; Yokoi & Diedrichsen, 2019), we detected sequence-specific activity patterns, i.e. activity patterns that differentiate between the tested motor sequences, in various cortical regions, even in session 1 (Fig. 6a). This allowed us to assess their reorganization across sessions.

**Figure 6.**
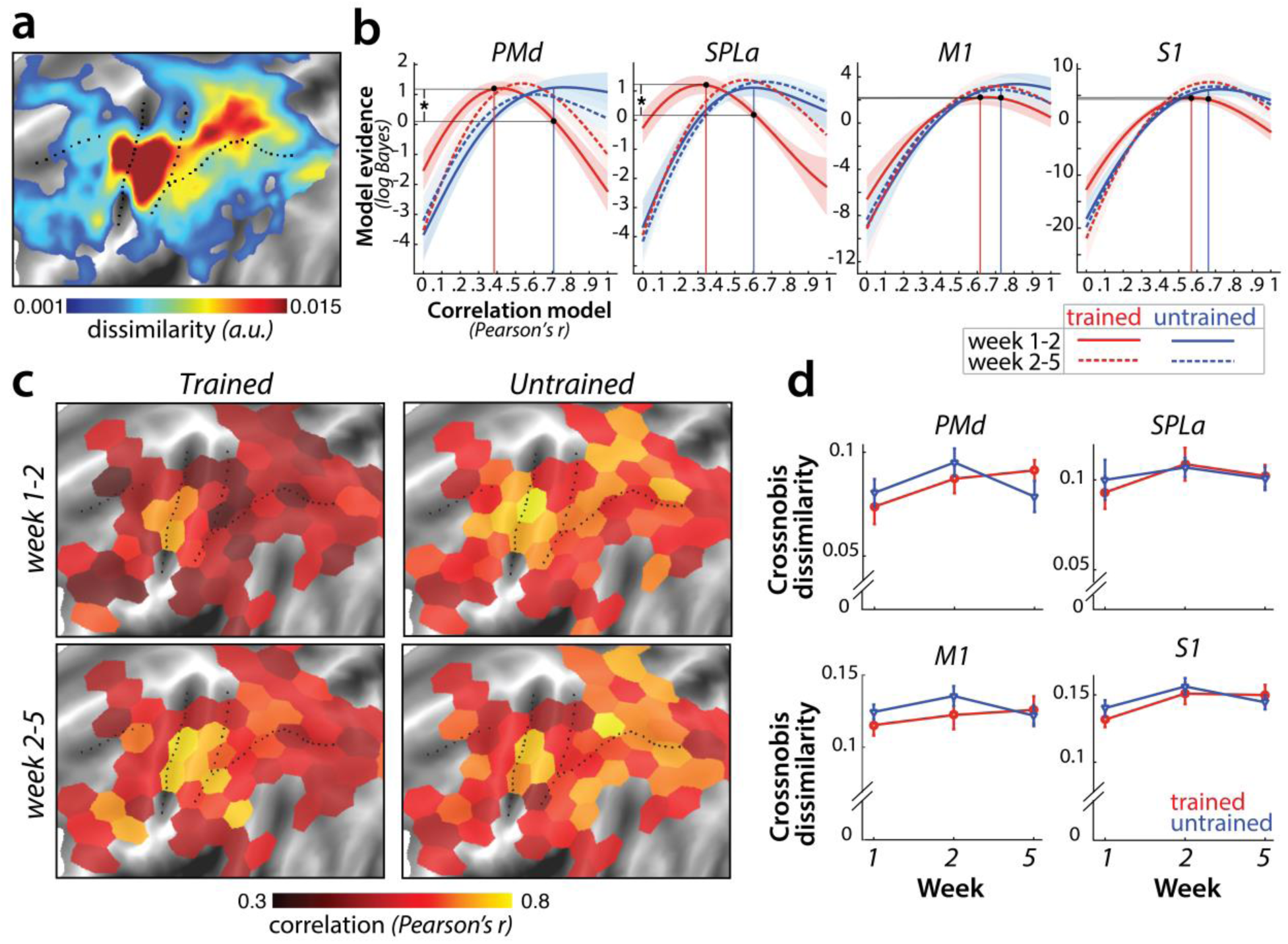
Sequence-specific activity patterns reorganize across sessions. **a)** Cortical surface map of crossnobis dissimilarities between activity patterns for different sequences in session 1. These regions encode which sequence is executed by the participant. **b)** Evidence explained by models of correlation values between *r* =0 and *r* =1 for sequence-specific patterns across weeks 1-2 (solid) and 2-5 (dashed), separately for trained (red) and untrained (blue) sequences. Evidence was assessed with a type-II log-likelihood, relative to the average log-likelihood across models. Shaded areas indicate standard error across participants. Difference between log-likelihoods can be interpreted as log-Bayes factor, with a difference of 1 indicating positive evidence. Horizontal lines are drawn for the winning correlation model for trained (red) and untrained (blue) patterns across weeks 1-2. Black dots are projections of the two winning models onto the correlation function of trained sequences across weeks 1-2. The horizontal lines from the two black dots indicate the likelihood of the trained data under the two models, which was tested in a crossvalidated t-test. **c)** Map displaying the correlation of the winning model for trained and untrained sequences across weeks 1-2 and 2-5. The correlation of the winning correlation model is shown in all tessels where the difference between evidence for winning model vs. worst-fitting model exceeds log-Bayes factor of 1 (averaged across participants). See **S6** for the difference in best model correlation between trained and untrained sequences, and an indication of tessels where the difference is significant, as based on the crossvalidated t-test. **d)** Crossnobis dissimilarities between trained and untrained sequence pairs across weeks. No significant effect of week, sequence type or their interaction was observed in any of the regions. Error bars indicate standard error across participants.

Our pre-registered hypothesis (https://osf.io/etnqc) was that earlier in learning sequence-specific activity patterns would change more for trained than untrained sequences, and would stabilize later in learning. In contrast to the other ideas tested in this paper, this was a novel hypothesis and not based on previous reports. Specifically, we predicted that the correlation of each sequence-specific pattern between weeks 1 and 2 should be lower for trained as compared to untrained sequences. The problem with performing a simple correlation analysis on the patterns, however, is that the estimated correlation will be biased by noise – i.e., more within-session variability for one set of sequences will result in a lower correlation (Diedrichsen, Yokoi, & Arbuckle, 2017). To address this problem, we used the pattern component modelling (PCM) framework which explicitly models and estimates the signal and noise for each session explicitly. Using this approach, we estimated the likelihood of each participants’ data under a series of models, each assuming a true correlation in the range between *0* (uncorrelated patterns) and *1* (perfect positive correlation; see Materials and Methods for details). Figure **6b** shows the log-likelihood for each specific correlation model relative to the mean across all models. In SPLa, the most likely correlation of the activity patterns for the trained sequences between weeks 1 and 2 was *r* =0.37. For week 2-5, the likelihood peaked at *r* =0.6. In contrast, the likelihood functions for untrained sequences indicated that the most likely model was between *r* =0.6-0.7 for both week 1-2 and 2-5. The advantage of this analysis is that we can be sure that the observed low correlation in week 1-2 for trained sequence was not due to increased noise. In fact, if the noise in one or both sessions was too high, then the model would be unable to distinguish between any of the correlation models – i.e. the likelihood curve would be a flat line.

To statistically assess the difference in correlations across trained and untrained sequences, we compared the likelihood of the data of trained sequences between two models: the best-fitting model for the trained sequences (*r* =0.37 in SPLa) and the correlation model best fitting the data of untrained sequences (*r* =0.6) (black dots and projections onto y-axis in Fig. 6b). To avoid double-dipping, the ‘best-fitting’ model was chosen on 25 participants (n-1) and the likelihood assessed on the left-out subject (see Materials and Methods). The difference in model evidence was significant for correlation between weeks 1-2 in SPLa (*t*_(25)_=2.88, *p*=8.0e-3). In contrast, no difference in correlation was observed later in learning, between weeks 2 and 5 (*t*_(25)_=1.21, *p*=0.24). A similar pattern of results was observed in PMd, with correlation of trained sequences significantly lower than that of untrained sequences between weeks 1 and 2 (*t*_(25)_=2.93, *p*=7.2e-3), but not between weeks 2 and 5 (*t*_(25)_=0.88, *p*=.39). No such change in correlation across weeks 1-2 was observed in M1 (*t*_(25)_=0.43, *p*=.67) or S1 (*t*_(25)_=1.72, *p*=0.097). Overall, we found significant evidence that sequence-specific trained patterns in SPLa and PMd reorganize more in weeks 1-2 as compared to the untrained sequences, and stabilize later on with learning, in line with our new pre-registered prediction.

To determine more generally where in the neocortex sequence-specific plasticity could be detected, we fit PCM correlation models to regularly tessellated regions spanning the cortical surface. Figure **6c** displays the correlation with the highest evidence for activity patterns across weeks 1-2 and 2-5; separately for trained and untrained sequences. In general, the highest correlations were found in core sensory-motor areas. Across weeks 1-2 for trained sequences, correlations were significantly lower in a number of dorsal premotor, inferior frontal, and parietal regions (Fig. 6c). Across the cortex, correlation for trained patterns increased for weeks 2-5, resulting in similar values which did not differ significantly between trained and untrained sequences for most tessels (see supplementary figure **S6**). Together, these results confirmed that sequence-specific activation patterns in secondary association areas show less stability early in learning, but stabilize later on.

Can we obtain further insight into *how* the sequence-specific patterns change in these areas? One specific preregistered prediction was that there would be an increase in distinctiveness (dissimilarity) between fMRI patterns underlying each trained sequence (Wiestler & Diedrichsen, 2013; Fig. 1e). To test this hypothesis, we calculated crossnobis dissimilarities (Walther et al., 2016) between sequence-specific activations, separately for trained and untrained sequences. In contrast to our prediction, no significant change in dissimilarity across weeks was observed in any of the predefined regions (Fig. 6d). This suggests that the reorganization observed for trained sequences early in learning did not increase the average distinctiveness of the sequence-specific patterns.

### Trained sequences elicit distinct patterns during full speed performance

In the last part of the experiment, we asked whether some of the negative findings (e.g. no changes in M1, no increase in dissimilarities for trained sequences) might have been due to the fact that participants were paced at a relatively slow speed. Matching the speed across sessions allows for the comparisons of changes in neural activity for exactly the same behavioral output (Karni et al., 1995; Lehéricy et al., 2005). However, it could be that controlling for speed impairs our ability to study brain representations of motor skill; simply because after learning, the system is not challenged enough to activate the neuronal representations supporting skilled performance. Consequently, several studies have not (Bassett et al., 2010; Wymbs & Grafton, 2015), or not strictly (Wiestler & Diedrichsen, 2013), matched performance across sessions or levels of training. To examine the effect of performance speed, we added a fourth scanning session *(fs)*, just a day after from the third session in week 5, in which participants were instructed to perform the sequence as fast as possible.

Performance during the 4th scan was 1010 ms faster than in the first session (*t*_(25)_=15.7, *p*=1.82e-14) and also 338 ms (*t*_(25)_=9.92, *p*=4.58e-10) faster for trained than for untrained sequences. Averaged over trained and untrained sequences, we found that the faster performance in this session led to an increase in activity across premotor and parietal areas (Fig. 7a,b). Although trained sequences were executed faster than untrained sequences, activation was still lower for trained compared to untrained sequences, similar to what we observed for paced performance (Fig. 7c; see S7a for statistical maps). In M1 and S1, we found no difference in activation between trained and untrained sequences (Fig. 7a; M1: *t*_(25)_=1.78, *p*=.09; S1: *t*_(25)_=1.69, *p*=.10). Overall, the pattern of results for evoked activation did not change qualitatively when participants performed at full speed.

**Figure 7.**
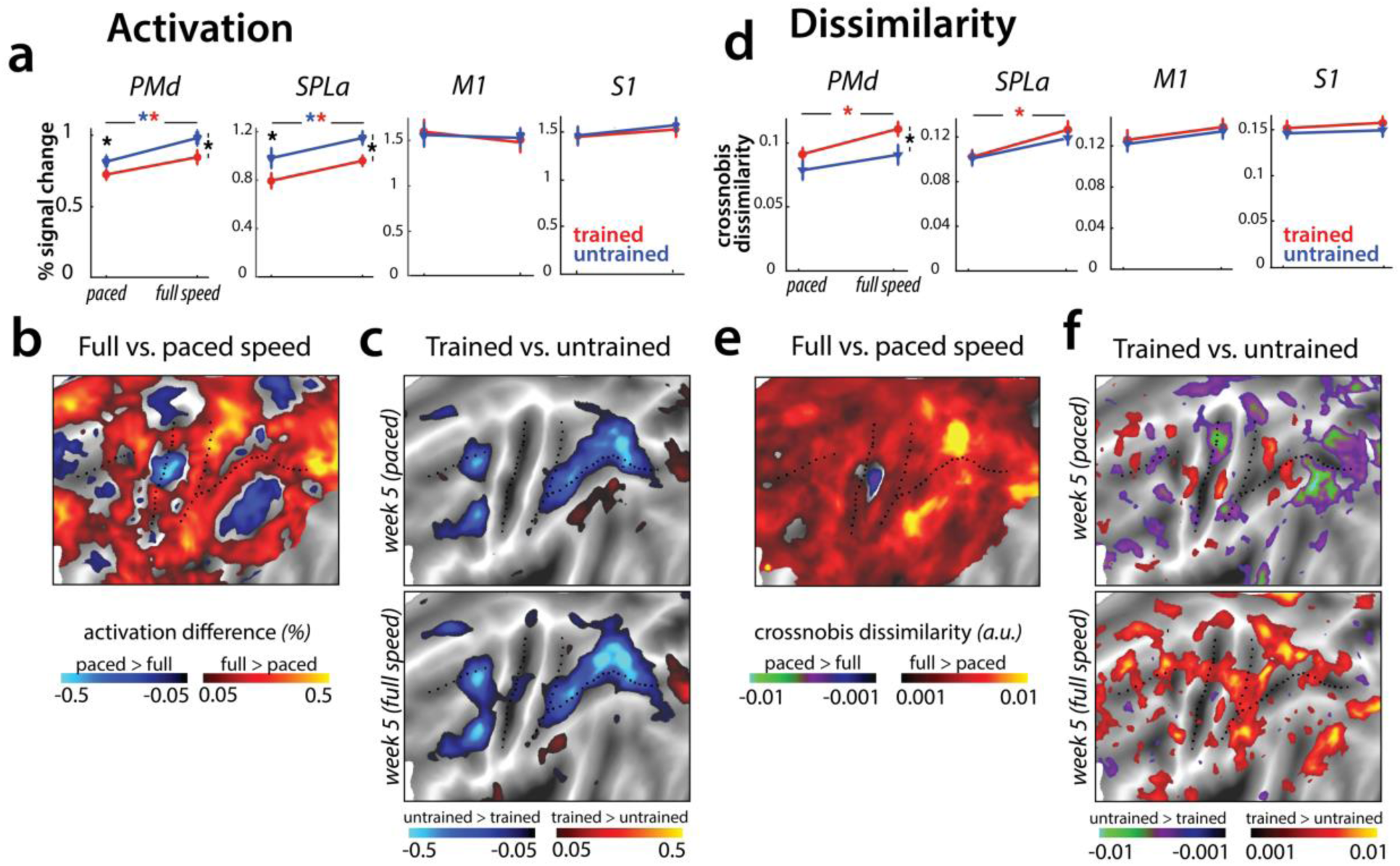
Speed-related changes in activation and dissimilarities. **a)** Overall activation in week 5 in paced and full speed sessions for trained (red) and untrained (blue) sequences. Activation was measured as percent signal change over resting baseline (* indicates *p*<.05). Error bars indicate standard error across participants. **b)** Increase in activation for full speed compared to paced speed in percent signal change, averaged across trained and untrained sequences. Red colors indicate an increase in activity during full speed performance compared to paced performance. Blue colors indicate higher activation during paced compared to full speed performance. **c)** Difference in activation elicited for trained relative to untrained sequences, during the paced and full speed sessions (see supplementary figure **S7a** for statistical maps). Trained>untrained is shown in red, untrained>trained in blue. **d)** Average crossnobis dissimilarity between sequence-specific patterns in paced and full speed sessions for trained and untrained sequences. Dissimilarities are significantly larger for trained (red), as compared to untrained (blue) patterns, in PMd for full-speed session (* indicates *p*<.05). Error bars indicate standard error across participants. **e)** Difference between crossnobis dissimilarities across full speed and paced sessions, averaged across trained and untrained sequences. Higher dissimilarities for full speed than paced session are shown in red, whereas blue/green hues indicate higher dissimilarities during paced than full speed session. **f)** Difference in dissimilarities for trained relative to untrained sequences, during the paced and full speed sessions. Trained>untrained is shown in red, untrained>trained in blue/green. Trained sequences elicited higher dissimilarities than untrained in full speed, but not paced session (see **S7b** for statistical *t*-maps).

Next, we examined whether the brain representations of individual sequences are similarly engaged at slow and fast speeds. The correlation between sequence-specific patterns was relatively high (*r*=0.62) across our regions of interest. We found no differences between the different regions (*F*_(3,75)_=1.47, *p*=.23), or sequence types (trained vs. untrained: *F*_(1,25)_=0.25, *p*=.62). Thus, the sequence-specific representations activated during performance at high skill level (full speed) are at least partly activated even when performance slowed down.

Having established that the mean activation results are replicated across paced and full-speed performance, and that similar sequence-specific representations are activated in both cases, we tested whether activation patterns for different trained sequences are more distinct during full speed performance, as reported in Wiestler & Diedrichsen (2013). Overall, crossnobis dissimilarities increased at full speed for trained sequences in PMd and SPLa (Fig. 7e). No such changes were found in M1 or S1. Moreover, trained sequences showed larger dissimilarities than untrained at full-speed performance across premotor and parietal cortices (Fig. 7f), which was not the case for the last paced session. In our predefined ROIs, this difference was significant for PMd (Fig. 7d), but also parietal areas showed significantly higher dissimilarities between trained sequences at full speed (supplementary figure **S7b**). This suggests that while activity patterns at full speed are correlated to those during paced performance, they are more distinguishable for trained sequences.

Could this effect be driven by behavioral performance, with trained sequences performed more differently at full speed (i.e. different speeds across trained sequences), while untrained sequences were performed at a more equal speed? To test for this, we calculated crossnobis dissimilarities between movement times associated with different trained and untrained sequences. The dissimilarities based on speed of performance did not differ significantly across trained and untrained sequences (*t*_(25)_=0.57, *p*=.57). Therefore, increased dissimilarity of trained compared to untrained patterns in premotor and parietal areas could not be explained by a difference in execution speed. Instead, this effect likely reflects changes in activity patterns underlying full speed skilled performance.

### Striatal activity patterns for trained sequences manifest at full speed performance

We observed learning-related changes in cortical association areas, but not in the primary motor cortex. Of course, learning could also be driven by neuronal changes in subcortical brain regions (Ashby, Turner, & Horvitz, 2010; Graybiel, 2016; Graybiel & Grafton, 2015; Hikosaka et al., 1999; Yin et al., 2009). The striatum in particular has been proposed as a structure where motor skills are stored (Kawai et al., 2015; Lehéricy et al., 2006). Inspecting changes in overall activity across sessions, we observed no difference in activity between trained and untrained sequences in either putamen or caudate nucleus (Fig. 8a).

**Figure 8.**
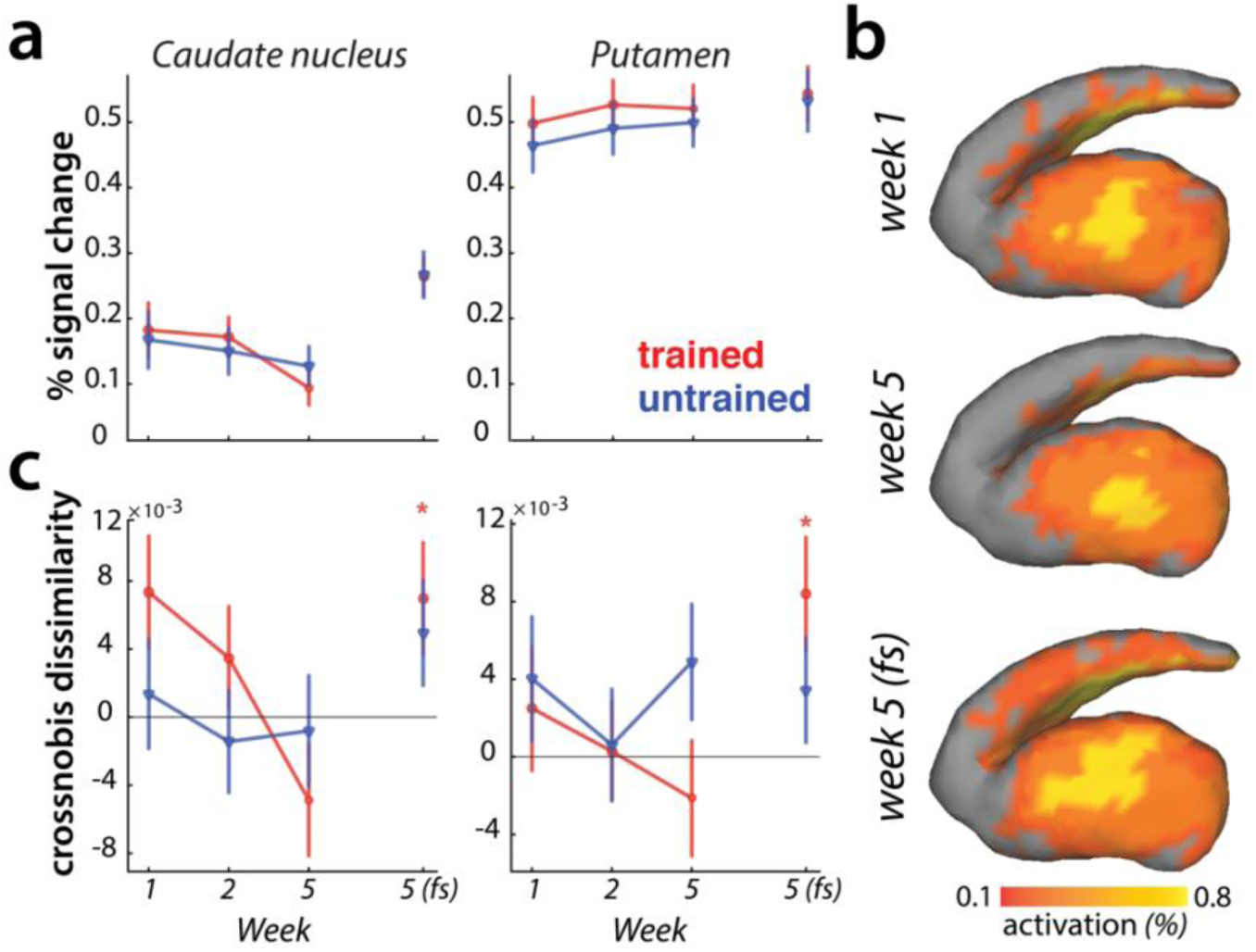
Striatal changes in activation and dissimilarities with learning. **a)** Overall activation (percent signal change over resting baseline), for trained (red) and untrained (blue) sequences. Activation did not differ across sessions, or sequence types in the striatum. Error bars indicate the standard error across participants. **b)** Activation during performance of trained sequences in the striatum across weeks 1, 5 (paced speed) and 5 (full speed – *fs*), averaged across sequences and participants. **c)** Crossnobis dissimilarities between activation patterns of sequence pairs, calculated separately for trained and untrained patterns. Dissimilarities were not significantly different for trained or untrained sequences during paced performance. At full speed, sequence-specific activity patterns amongst trained sequences differed significantly in both caudate nucleus and pallidum (* indicates *p*<.05). Error bars indicate the standard error across participants.

Previous experiments have reported that with learning, activation moves from more ‘cognitive’ areas of the striatum (i.e. caudate nucleus) to more ‘motor’ areas (i.e. putamen) (Coynel et al., 2010; Lehéricy et al., 2005; Reithler, van Mier, & Goebel, 2010). Our data fails to replicate this result: Both the visual inspection (Fig. 8b), and statistical quantification of the mean pattern difference for trained and untrained sequences across sessions revealed no such learning-specific shift of mean striatal activation pattern with learning.

Lastly, we examined if the striatum represents individual sequences. During the paced sessions, activity patterns for different sequences were not distinguishable in either caudate nucleus or putamen (Fig. 8c). However, during full speed performance trained sequences elicited distinct activity patterns in both regions (i.e. crossnobis dissimilarity>0: caudate nucleus: *t*_(25)_=2.27, *p*=0.032; putamen: *t*_(25)_=2.44, *p*=.022; Fig. 8c). This effect was specific to the trained sequences, with untrained sequences still exhibiting undistinguishable patterns of activity at full speed. Thus, we found some evidence that trained motor sequences are represented in the form of distinct activity patterns in the striatum during full speed skilled performance.

## Discussion

Here we present a large longitudinal motor sequence learning study that allowed us to systematically investigate several previously proposed fMRI signatures of motor learning, including one new hypothesis concerning the change in multivariate activity patterns with learning. The existing literature, with its diversity of experimental protocols and analysis approaches, does currently not provide a consistent picture of learning-related changes. This inconsistency is exacerbated by the fact that most papers prioritize making new claims over re-examining previously established findings. Consequently, it is very hard to assess the replicability of most past findings. We address this issue here by a) producing a well-powered, longitudinal data set that tackles some of the methodological inconsistencies (i.e. speed matching), b) pre-registering both design and hypotheses, and c) making data and analysis pipelines openly available, such that other hypotheses and analyses techniques can be freely tested.

Our findings reveal that parietal and premotor areas show widespread decreases in overall activation, as well as reorganization of sequence-specific patterns early in learning. Additionally, we observed that sequence specific patterns in these areas (as well as the striatum) were more distinct during full speed performance. In contrast to this set of results, none of these learning-specific metrics were detected in M1, even after 5 weeks of training.

On the one hand, our lack of any observable change in M1 activation contradicts some prior results, where increased activation in M1 was observed for matched performance after learning (Karni et al., 1995; Matsuzaka, Picard, & Strick, 2007; Penhune & Doyon, 2002; Steele & Penhune, 2010; Vahdat et al., 2015), and does not align with reports of M1 stimulations influencing consolidation or storage of motor skills (in motor sequence tasks: Kang & Paik, 2011; Nitsche et al., 2003; Reis et al., 2009; Waters-metenier, Husain, & Wiestler, 2014; in other motor tasks: Classen, Liepert, Wise, Hallett, & Cohen, 1998; Galea, Vazquez, Pasricha, Orban De Xivry, & Celnik, 2011; Hadipour-Niktarash, Lee, Desmond, & Shadmehr, 2007). We also found no support for a combination of increases and decreases of activation with training, which would lead to an overall change of the mean activity pattern (Steele & Penhune, 2010).

Instead, our results suggest that the pattern of neural activity in M1 does not change as participants become more skilled at producing motor sequences. This is consistent with a recent line of evidence demonstrating that M1 does not change activation with learning (Huang et al., 2013), and primarily encodes single movement elements, rather than sequences (Yokoi, Arbuckle, & Diedrichsen, 2018; Russo et al., 2019). Somewhat more surprisingly, we also observed no difference in overall M1 activation during full speed performance, when performance was considerably faster for trained sequences. This suggests that the activity increases related to faster movement speeds are compensated for by the shorter duration spent on the task.

Primary somatosensory cortex in many ways paralleled the results observed in M1. We observed no overall activation change, or change in the sequence-specific pattern correlation across sessions. The only exception was the observed shift in the mean activation pattern across sessions. One possible explanation is that feedback-related sensory activity in S1 undergoes some plastic changes with learning. This is consistent with a recent study demonstrating that S1, but not M1, is involved during consolidation of motor skills (Kumar, Manning, & Ostry, 2019; for a review on somatosensory plasticity in motor learning see Ostry & Gribble, 2016).

In contrast to the limited evidence of learning-related changes in primary somatosensory and primary motor areas, higher order association areas (e.g. parietal and premotor cortices) displayed an array of learning-related changes. First, activation decreased in areas involved in sequence execution, with larger decreases for trained as compared to untrained sequences. This result contrasts with other previous studies reporting increases in activation in premotor areas with learning (Grafton et al., 2002; Honda et al., 1998; Penhune & Doyon, 2002; Vahdat et al., 2015). Partially responsible for these inconsistencies may be a publication bias, favoring reports of signal increases over signal decreases with learning. For example, a recent metanalysis reanalyzed evidence for signal increases in the main text, while moving the (matched) evidence for signal decreases into the supplementary materials (Hardwick et al., 2013). Our data corroborates a number of recent studies reporting reduced activation in task-evoked premotor and parietal areas (Steele & Penhune, 2010; Wiestler & Diedrichsen, 2013; Wu et al., 2004).

The only activation increases for trained relative to untrained sequences were observed in areas that were suppressed below baseline during sequence execution. This has also been previously reported in a motor sequence learning study (Tamás Kincses et al., 2008), where deactivation was larger during performance of trained than random sequences. These areas include the precuneous, temporal parietal junction and the cingulate, regions commonly assigned to the default mode network (Raichle et al., 2001; Shulman et al., 1997). This group of regions is more activated during rest than during task performance, and has been associated with functions such as episodic memory retrieval and attention to internal states (Andrews-Hanna, Reidler, Sepulcre, Poulin, & Buckner, 2010; Gusnard, Akbudak, Shulman, & Raichle, 2001). Our observation of decreased inhibition of the default mode network likely reflects central attentional resources being freed up, allowing participants to engage in other mental processes (e.g., daydreaming) while performing the task. Thus, this release from initial deactivation is possibly task-irrelevant, reflecting increased automaticity with learning (Shamloo & Helie, 2016).

Overall, changes in average activation are relatively hard to interpret, as they could reflect a combination of numerous factors. As a more direct fMRI metric of plasticity, we suggest to inspect changes in the sequence-specific activity patterns, since these constitute a more likely fMRI correlate of the sequences-specific performance advantage observed after training. In this project, this provided us with two key insights of how activation patterns reorganize in association areas with learning. First, activity patterns associated with each of individual trained sequences, changed to a greater extent earlier in learning, and stabilized later. This finding resonates with several animal studies suggesting that the emergence of skilled behavior is associated with early plasticity and later stabilization of neuronal activity patterns (Makino et al., 2017; Peters et al., 2017). Here we report a similar effect in humans, and advance these findings by demonstrating that this reorganization occurs at the level of sequence-specific patterns. In past studies using rodent models, sequence-specific patterns could not be dissociated from the overall activity pattern, as the animals were only trained on production of a single sequence. Additionally, by pacing participants’ speed, we were able to cleanly dissociate changes in the organization of activity patterns from changes in the behavioral performance or variability. Second, activation patterns became more distinct for trained sequences at full speed. This indicates that the engagement of specific neuronal subpopulations for different sequences is particularly important when pushing the limit of performance.

While our study focused on the role of cortical areas in motor sequence learning, we also examined activation in the striatum, which has been suggested to play a critical role in skilled performance (Graybiel & Grafton, 2015; Kawai et al., 2015; Otchy et al., 2015). In contrast to previous fMRI studies (Coynel et al., 2010; Lehéricy et al., 2006; Reithler, van Mier, & Goebel, 2010), we did not find clear evidence for differences in overall activity, or shifts of the overall activity pattern with learning. Nonetheless, we observed distinguishable striatal activation patterns for different trained sequences at full speed, in line with a recent report showing distinguishable striatal patterns for performance of consolidated motor sequences (Pinsard et al., 2018). While by itself the finding of differential sequence-specific activity patterns is not evidence for a causal role of the striatum in the production of skilled behaviors, it is a necessary condition for such a functional role. Therefore, our results here are in line with the proposed involvement of the striatum in motor sequence learning.

An important feature of our design was that we collected imaging data in the trained state, both when performance was clamped to the initial speed, and when participants performed as fast as possible. Previous studies have usually included only one of these two options, making direct comparisons difficult (see Lutz et al., 2004 for an examination of various execution speeds on BOLD activity and Orban et al., 2010 in a motor learning context). Our results provide two important insights: first, in terms of the overall fMRI activation, the pattern of results remained the same for paced vs. full speed performance. This indicates that, in this specific case, the increased motor demands and the decreased time on task averaged out. In general, however, these two factors may not balance perfectly – therefore paced performance may be a better choice when comparing overall activation across sessions. Second, even though slow and paced performance in the trained state activated sequence-specific activation patterns, these were much stronger when performing at maximal speeds. Thus, for questions regarding the fine-grained patterns, it might be more suitable to challenge the system fully.

Of course, our list of inspected fMRI metrics of learning was not exhaustive. For instance, we did not investigate whether various fMRI correlates of learning predict behavioral outcomes, or how functional connectivity and network metrics change with learning, partly because of the absence of specific predictions. Pre-registration of hypothesis are especially important for these analyses, since the search space of possible tests becomes exponentially larger (e.g. correlating all possible brain metrics with all possible behavioral metrics; or using various metrics to assess inter-regional relationships). However, we hope that our dataset, upon its public release, can serve as a resource for other researchers to (re-)test novel predictions about learning related changes.

## Conclusion

The search for neural substrates of learning is a daunting task: the acquisition of longitudinal data sets is work intensive, and the large dimensionality of possible brain metrics makes the search difficult (Poldrack, 2000). Historically, the question was simplified by studying activation increases in single areas as proxies for motor ‘engram’ localization (Berlot, Popp, & Diedrichsen, 2018). Here we found no evidence for such activation increases; instead we observed widespread and distributed decreases in activation across cortical areas. In contrast, subtler changes in the distributed patterns of fMRI activity have the potential to provide more direct metrics of plasticity. Increased pattern reorganization (across weeks), and larger pattern separation for trained sequences was found across prefrontal, parietal, and striatal regions. These metrics may be useful as general fMRI correlates of neural reorganization beyond the domain of motor learning.

## Materials and Methods

### Participants

Twenty-seven volunteers participated in the experiment. One of them was excluded because field map acquisition was distorted in one of the four scans. The average age of the remaining 26 participants was 22.2 years (SD = 3.3 years), and the sample included 17 women and 9 men. All participants were right-handed and had no prior history of psychiatric or neurological disorders. They provided written informed consent to all procedures and data usage before the study started. The experimental procedures were approved by the Ethics Committee at Western University.

### Apparatus

Participants performed finger sequences with their right hand on an MRI-compatible keyboard (Fig. 2a), with keys numbered 1-5 for thumb-little finger. The keys had a groove for each fingertip and were not depressible. The force of isometric finger presses was measured by the force transducers (FSG-15N1A, Sensing and Control, Honeywell; dynamic range 0-25 N) mounted underneath each key with an update rate of 2 ms. A key press was recognized when the sensor force exceeded 1 N. The measured signal was amplified and sampled at 200 Hz.

### Learning paradigm

Participants were trained to execute six 9-digit finger sequences over a period of five weeks (Fig. 2a). They were split into two groups with trained sequences of one group constituting the untrained sequences for the other group and vice versa. Finger sequences of both groups were matched as closely as possible in terms of the starting finger, number of finger repetitions in a sequence and first-order finger transitions. This counterbalancing between the groups ensured that any of the observed results were not specific to a set of chosen trained sequences.

In the pre-training session prior to the first scan (Fig. 2b), participants were acquainted with the apparatus and task performed during scanning. Sequences executed during this pre-training session were not encountered later on in the experiment.

During the training sessions, participants were trained to perform the six sequences as fast as possible. They received visual feedback for the correctness of their presses with digits turning green for a correct finger press and red for an incorrect one. After each trial, participants received points based on the accuracy and their movement time (MT – time from the first press until the last finger release in the sequence; Fig. 2c). Trials executed correctly and faster than participant’s median MT from the previous blocks were rewarded with 1 point. If participants performed correctly and 20% faster than the median MT from previous blocks, they received 3 points. If they made a mistake or performed below their median MT, they received 0 points. Participants performed each sequence twice in a row: digits were written on the screen for the first execution, but removed for the second execution so that participants had to perform the finger sequence from memory. Training sessions were broken into several blocks, each consisting of 24 trials (4 trials per trained sequence), with time between blocks to rest. At the end of each block, participants received feedback on their error rate, median MT and points obtained during the block. If participants performed with an error of <15% and faster than the previous median MT, the MT threshold was updated. This design feature was chosen to maintain participants’ motivation to execute the sequences as fast as possible, within the allowed error range.

During the behavioral test sessions (Fig. 2d), participants executed sequences they were trained on as well as untrained matched sequences, which were randomly interspersed. All sequences were still performed twice in a row, with numbers on the screen present on both executions.

### Experimental design during scanning

Participants underwent four scanning sessions (Fig. 2d) – with the first one before learning regime started, the second after a week and two more scans after completion of the 5 training weeks. Each scanning session consisted of eight functional runs. We employed an event-related design, randomly intermixing execution of trained and untrained sequences. Each sequence was repeated twice in a row (with digits always present on the screen), and there was a total of six repetitions per sequence in every run. Each trial started with 1 second preparation time, during which the sequence was presented on the screen. After that time, a ‘go’ signal was displayed as short pink line underneath the sequence numbers. In scanning sessions 1-3, this line started expanding below the written numbers, indicating the speed at which participants were required to press along. In scanning session 4, only a short line was presented in front and underneath the sequences. When the line disappeared, this signaled a ‘go’ cue for participants to execute the presented sequence as fast as possible. The execution phase including the feedback on overall performance lasted for 3.5 seconds, and the inter-trial interval was 0.5 seconds (see supplementary figure **S2**). Each trial lasted for 5 seconds. Five periods of rest, each 10 seconds long, were added randomly between trials in each run to provide a better estimate of baseline activation.

### Image acquisition

Data was acquired on a 7-Tesla Siemens Magnetom scanner with a 32-receive channel head coil (8-channel parallel transmit). Anatomical T1-weighted scan was acquired at the beginning of the first scanning session, using a magnetization-prepared rapid gradient echo sequence (MPRAGE) with voxel size of 0.75×0.75×0.75 mm isotropic (field of view = 208 × 157 × 110 mm [A-P; R-L; F-H], encoding direction coronal). Functional data were acquired using a sequence (GRAPPA 3, multi-band acceleration factor 2, repetition time [TR] = 1.0 s, echo time [TE] = 20 ms, flip angle [FA] = 30 deg). We acquired 44 slices with isotropic voxel size of 2×2×2 mm. For estimating magnetic field inhomogeneities, we additionally acquired a gradient echo field map. Acquisition was in the transversal orientation with field of view 210 × 210 × 160 mm and 64 slices with 2.5 mm thickness (TR = 475 ms, TE = 4.08 ms, FA = 35 deg).

### First-level analysis

Functional data were analyzed using SPM12 and custom written MATLAB code. Functional runs were corrected for geometric distortions using fieldmap data (Hutton et al., 2002), and head movements during the scan (3 translations: x, y, z; 3 rotations: pitch, roll, yaw), and aligned across sessions to the first run of the first session. The functional data were then co-registered to the anatomical scan. No smoothing or normalization to an atlas template was performed.

Preprocessed data were analyzed using a general linear model (GLM; Friston et al., 1994). Each of the performed sequences was defined as a separate regressor per imaging run, resulting in 12 regressors per run (6 trained, 6 untrained sequences), together with intercept for each of the functional runs. The regressor was a boxcar function starting at the beginning of the trial and lasting for trial duration. The boxcar function was convolved with a hemodynamic response function, with a time to peak of 5.5 seconds, and a manually adjusted onset to best fit each participant’s average evoked response. This analysis resulted in one activation estimate (beta image) for each of the 12 conditions per run, in each scanning session.

### Surface reconstruction and regions of interest

We reconstructed individual subjects’ cortical surfaces using FreeSurfer (Dale, Fischl, & Sereno, 1999). All individual surfaces were aligned to the FreeSurfer’s Left-Right symmetric template (workbench, 164k nodes) via spherical registration. To detect sequence representation across the cortical surface, we used a surface-based searchlight approach (Oosterhof, Wiestler, Downing, & Diedrichsen, 2011), where for each node we selected a circular region of 120 voxels in the grey matter. The resulting analyses (dissimilarities between sequence-specific activity patterns, see below) was assigned to the center node. As a slightly coarser alternative to searchlights, we performed regular tessellation of cortical surface into 162 tessels per hemisphere. This allowed us to fit correlation models (see below) across the cortical surface, while not being as computationally intensive as searchlight analyses.

We defined four regions of interest to cover primary somato-motor regions as well as secondary associative regions. M1 was defined using probabilistic cytoarchitectonic map (Fischl et al., 2008) by including nodes with the highest probability of belonging to Brodmann area (BA) 4, while excluding nodes more than 2.5 cm from the hand knob (Yousry et al., 1997). Similarly, S1 was defined as nodes related to hand representation in BA 1, 2 and 3. Additionally, we included dorsal premotor cortex (PMd) as the lateral part of the middle frontal gyrus. The anterior part of the superior parietal lobule (SPLa) was defined to include anterior, medial and ventral intraparietal sulcus. We also defined caudate nucleus and putamen as striatal regions of interest. The definition was carried out in each subject using FSL’s subcortical segmentation.

### Changes in overall activation

We calculated the average percent signal change for trained and untrained sequences (averaged across the 6 trained and 6 untrained sequences) relative to the baseline for each voxel. The resulting volume map was projected to the surface for each subject, and a group statistical *t*-map was generated across subjects. Statistical maps were thresholded at *p*<.01, uncorrected, and the family-wise error corrected *p*-value for the size of the peak activation and activation cluster size was determined using a permutation test. Specifically, we ran 1000 simulations where we randomly flipped the sign of the contrast for subjects (chosen at random out of 226 possible permutations). The rationale behind this is that under the null hypothesis, there should be no difference between the two conditions, and the sign of each contrast should be interchangeable. As for the data, we thresholded the statistical map from each permutation, and recorded the peak *t*-value (across the map) and the size of the largest cluster. The real data was then compared against this distribution to assess the probability of the observed *t*-value and cluster-size under the null hypothesis.

Additionally, we assessed changes in percent signal in predefined regions of interest (M1, S1, PMd, SPLa). This was performed in the native volume space of each subject. To do so, we averaged the percent signal change of voxels belonged to a defined region per subject and quantified activation changes across subjects using ANOVAs and t-tests across subjects.

Besides overall activation, we also examined *relative* changes in elicited activation for trained sequences across sessions. This was done by normalizing (z-scoring) the percent signal change surface maps across voxels, separately for each subject. Normalization was applied both map-wise (for Fig. 5b), as well as for each of the pre-defined ROIs separately (for cross-sections in Fig. 5c). Statistical assessment of the difference between relative evoked activation pattern for trained vs. untrained sequence was carried out by calculating cosine angle dissimilarities between the mean evoked patterns. Cosine angle dissimilarity was chosen because it is not sensitive to overall magnitude in activation, and therefore assesses the difference in the relative activation distribution.

### Dissimilarities between sequence-specific activity patterns

To evaluate which cortical areas display sequence-specific encoding, we performed a searchlight analysis calculating the dissimilarities between evoked beta patterns of individual sequences. Beta patterns were first multivariately prewhitened (standardized by voxels’ residuals and weighted by the voxel covariance matrix), which has been found to increase the reliability of dissimilarity estimates (Walther et al., 2016). We then calculated the cross-validated squared Mahalanobis dissimilarities (i.e. crossnobis dissimilarities) between evoked sequence patterns (66 dissimilarity pairs for 6 trained and 6 untrained sequences). These dissimilarities were then averaged overall, as well as separately for pairs within trained sequences, and within untrained sequences. This metric was used both for searchlight analysis and calculation of metric within predefined regions (cortical and striatal). The cortex surface maps contrasting dissimilarities between trained and untrained sequences were corrected for multiple comparisons using permutations, as described above for percent signal change surface maps.

### Pattern component analyses: modelling sequence-specific correlation across sessions

Correspondence of sequence-specific patterns across sessions was quantified using pattern component modelling (PCM; Diedrichsen et al., 2017). This framework is superior at estimating correlations than simply performing Pearson’s correlation on raw activity patterns, or even in a crossvalidated fashion. The main problem with estimating correlations on data is that activation patterns are biased by noise, which varies across scanning sessions, and would therefore underestimate the true correlation. PCM separately models the noise and signal component, and can in this way combat the issue more than simply performing crossvalidation would. We designed 30 correlation models with correlations between 0 and 1 in equal step sizes and assessed the group likelihood of the observed data under each model.

Subsequent group inferences were performed using crossvalidated approach on assessing individual log-Bayes factors (model evidence). A crossvalidated approach was used to ensure that our choice of ‘best-fitting models’ and the evidence associated was independent and did not involve double-dipping. Specifically, we used n-1 subjects to determine the best-fitting models for trained and untrained patterns and recorded the log-Bayes factors for those two correlation models on the left-out subject. This was repeated across all subjects and a t-test was performed on the recorded log-Bayes factors (i.e. out-of-sample model evidences). The same evaluation was performed for pre-defined regions of interest (Fig. 6b), as well as a regular tessellation across the cortical surface (Fig. 6c).

## Acknowledgements

The work was supported by an Ontario Trillium Scholarship to EB, an NSERC Discovery Grant (RGPIN-2016-04890) to JD, and the Canada First Research Excellence Fund (BrainsCAN). We thank Giacomo Ariani, Tamar Makin, Andrew Pruszynski, and Atsushi Yokoi for helpful comments on the manuscript.

## Author contributions

EB, NJP and JD designed the experiment; EB and NJP programmed the experiment; EB and NJP collected the data; EB analyzed the data; EB prepared figures; EB drafted manuscript; EB, NJP and JD edited and revised the manuscript.

## Competing interests

The authors declare no competing interests.

## Supplementary Figures

**Figure S2.**
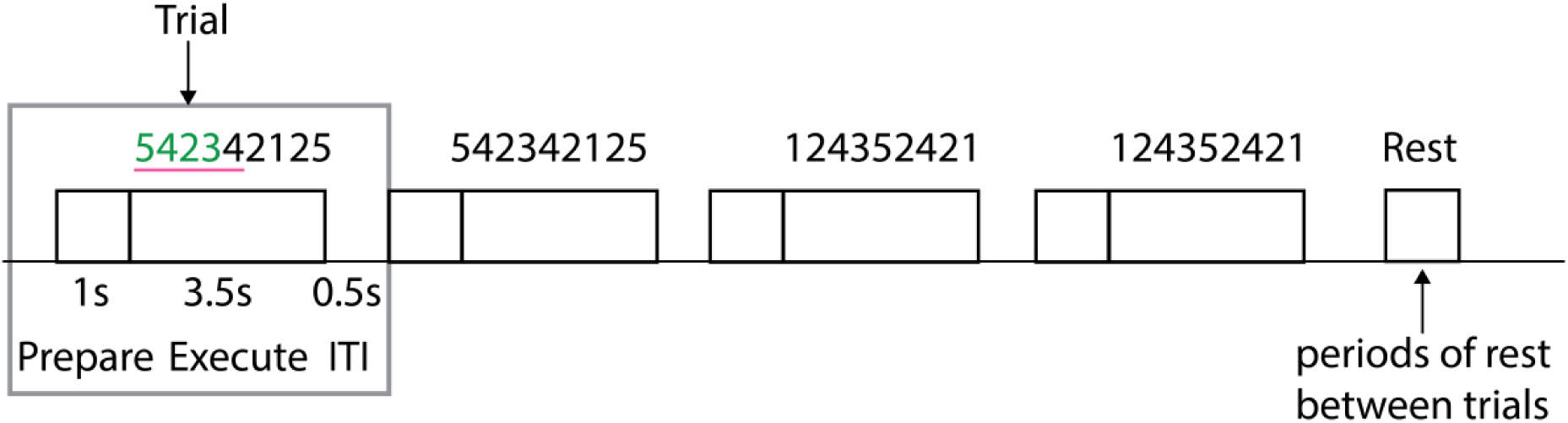
Experimental trial structure during scanning sessions. Each trial consisted of a preparation period, execution period and inter-trial-interval (ITI), during which the feedback was presented on correctness of the trial. Each sequence was presented twice in a row. Periods of rest were added in-between the trials.

**Figure S4.**
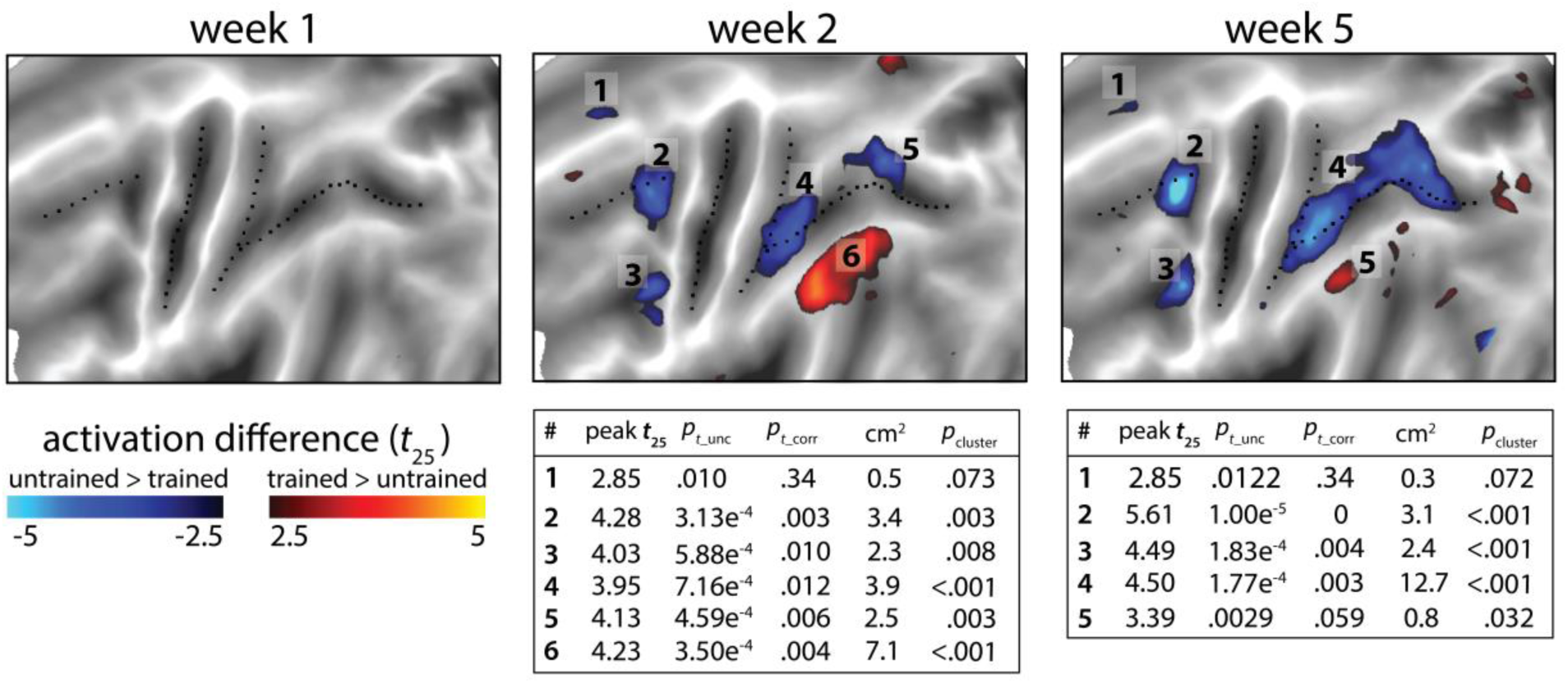
Statistical maps for the trained vs. untrained contrasts on elicited activation in each session. Trained>untrained is shown in red, untrained>trained in blue. Maps were thresholded at a *t*_25_=±2.5, *p*<.01 uncorrected for a two-sided *t*-test. Tables show peak *t*-value and size (in cm2) for each super-threshold cluster (indicated by numbers) for maps of week 2 and 5. *pt*_unc is the uncorrected *p*-value for the peak of each cluster. Family-wise error corrected *p*-values were determined using permutation testing for the peak *t*-value (*pt*_corr) and cluster size (*p*cluster).

**Figure S6.**
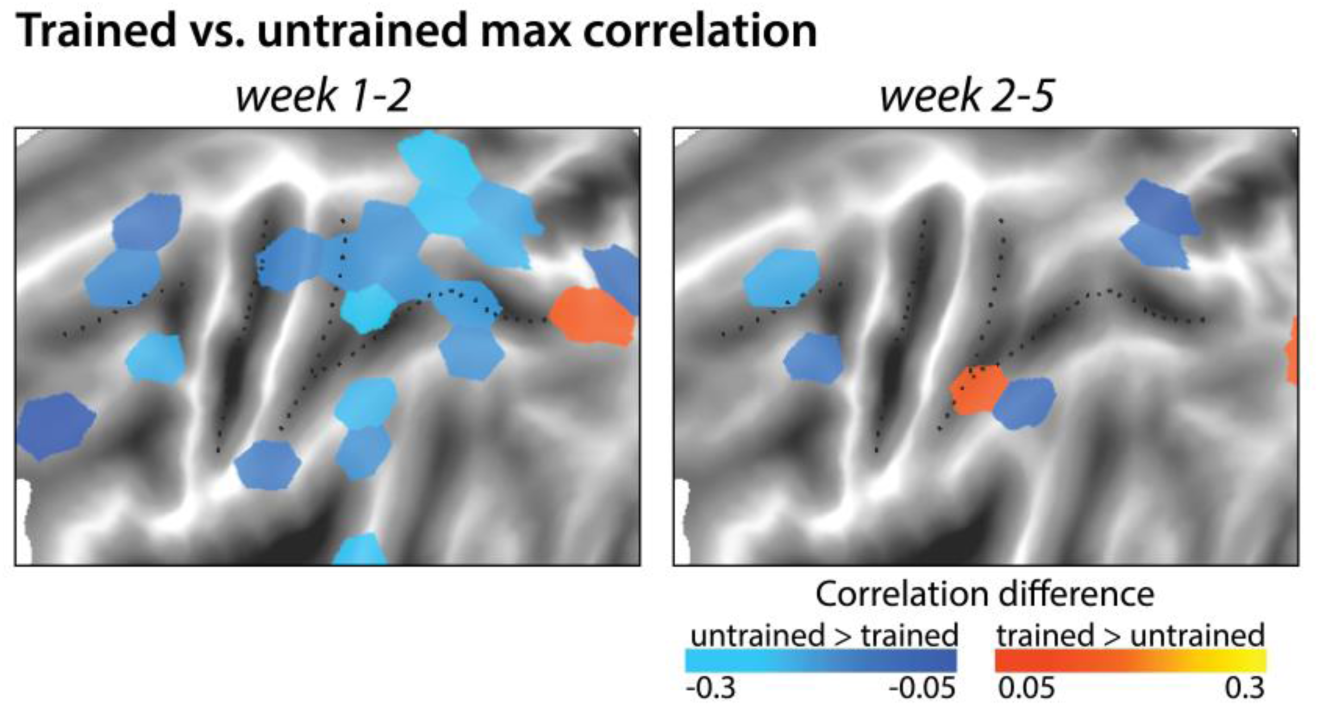
Difference between correlation of winning model for trained and untrained sequences. Difference between the correlations of the winner models for trained and untrained sequences, separately for week 1-2 and week 2-5. Blue indicates a lower correlation across weeks for trained than untrained patterns of activity. The correlation difference values are plotted in tessels where the difference in model evidence was significant, as based on the cross-validated *t*-test (for two-sided *p*<.05).

**Figure S7.**
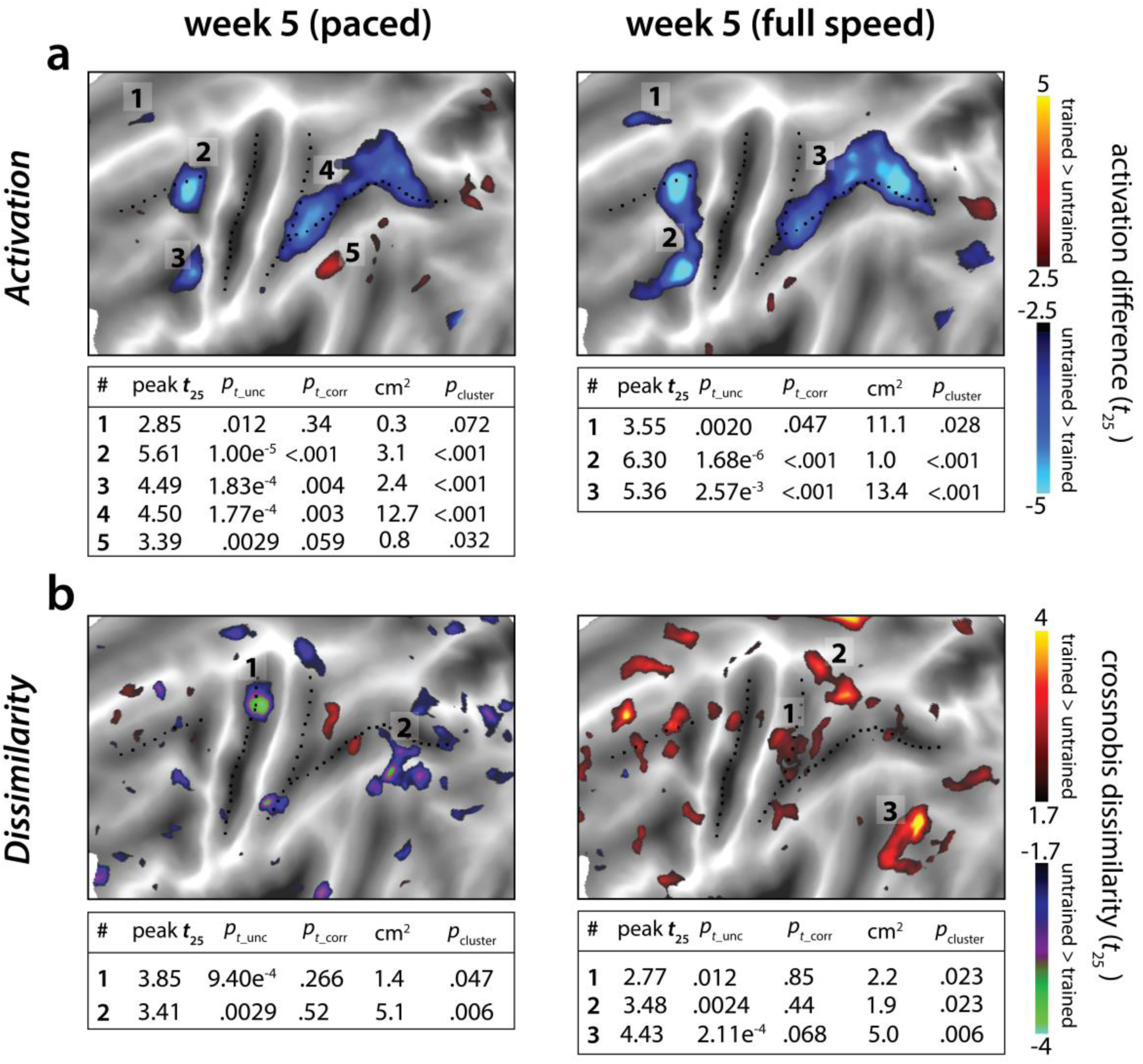
Statistical maps for trained vs. untrained contrasts in week 5 (paced) and 5* (full speed) sessions. Trained>untrained is shown in red, untrained>trained in blue. **a)** Statistical contrast for average activation. Maps were thresholded at a *t*_25_=± 2.5, *p*<.01 uncorrected for a two-sided *t*-test. Tables show peak *t*-value and size (in cm2) for each super-threshold cluster. *pt*_unc is the uncorrected p-value for the peak of each cluster. Family-wise error corrected *p*-values were determined using permutation testing for the peak *t*-value (*pt*_corr) and cluster size (*p*cluster). **b)** Statistical contrast for average dissimilarity of sequence-specific activity pattern. Map was thresholded at *t*_25_=± 1.7, *p*<.05, uncorrected. Statistical quantification using permutation tests is in the table below each map.

